# Thalamus-derived glutamate is required for early specification of layer 4 neurons in the sensory cortex

**DOI:** 10.64898/2026.05.26.727990

**Authors:** Daven Rock, Naomi Stow, Cindy Yu, Yasushi Nakagawa

**Affiliations:** Department of Neuroscience, University of Minnesota Medical School. Minneapolis, MN 55455 USA

## Abstract

Elucidating the mechanisms that control the formation of the mammalian neocortex is crucial for understanding brain functions. Synaptic activity of thalamocortical axons (TCAs), mediated by glutamate, exerts a major extrinsic influence on the maturation of their target layer 4 neurons in postnatal primary sensory cortex. However, TCAs reach the sensory cortex during mid-embryonic stages in mice, when neurons of future superficial layers, including layer 4, are still being generated from radial glia (RGs) or intermediate progenitor cells (IPCs), well before the formation of direct synapses. We previously showed that TCAs are required for the production and specification of the proper number of layer 4 neurons in sensory areas, and that part of these area-specific roles is played by the thalamus-derived molecule VGF. However, the role of TCA-derived glutamate prior to synapse formation has remained unclear. In this study, we used mutant mice lacking *vGluT2*, a vesicular glutamate transporter expressed in the embryonic thalamus, and found that vesicular release of thalamus-derived glutamate is required for the proper production and specification of layer 4 neurons in the sensory cortex by the neonatal stage, through mechanism distinct from those involving VGF. Our findings reveal that multiple molecular cues produced by incoming TCAs play distinct roles in the production and specification of layer 4 neurons in the sensory cortex.

## Introduction

The adult neocortex is composed of both excitatory and inhibitory neurons that are organized along radial (layers) and tangential (areas) axes. Among the many functionally distinct neocortical areas, primary sensory areas possess a well-developed layer 4, which serves as the main recipient layer of thalamocortical axons (TCAs) conveying first-order information for vision, somatic sensation and hearing. Decades of studies have addressed the roles of TCAs in the many aspects of neocortical development (Nakagawa 2026). Synaptic activity of TCAs in the postnatal mouse cortex has been shown to be crucial for the morphological and functional maturation of layer 4 neurons in the sensory cortex (Callaway & Borrell 2011, Li et al 2013, Narboux-Neme et al 2012, Pouchelon et al 2014). Because TCAs arrive in the embryonic cortex during the period of superficial layer neurogenesis, we recently investigated the early roles of TCAs prior to synapse formation using thalamus-specific *Gbx2* conditional knockout (cKO) mice, which lack most TCAs in the cortex (Monko et al 2022). In these mice, the number of cells produced at embryonic day 14.5 (E14.5) was reduced in the sensory cortex, but not in the motor or prefrontal cortex. In addition, the fate of neurons generated at E14.5 was shifted from layer 4 to layer 2/3. This shift was observed as early as postnatal day 1 (P1), before the formation of direct synapse between TCAs and prospective layer 4 neurons. These findings provide direct evidence that TCAs have a previously unrecognized role in regulating progenitor proliferation and early neuronal fate specification in the sensory cortex.

However, the molecular mediators underlying these roles are only beginning to be understood. *Vgf* (non-acronymic) encodes a precursor protein that is cleaved into more than a dozen neuropeptides (Bartolomucci et al 2011, Wang et al 2022), and is expressed in sensory thalamic nuclei as early as E12.5 in mice (Monko et al 2022). Thalamus-specific deletion of *Vgf* results in a shift in the fate of E14.5-born neurons from layer 4 to layer 2/3, similar to what is observed in *Gbx2* cKO mice; however, these mutants do not show a reduction in overall neuronal production (Monko et al 2022).

In addition to *Vgf*, *vGluT2*, which encodes a vesicular glutamate transporter, is also expressed in the early embryonic thalamus (Allen Developing Mouse Brain Atlas). Glutamate is an interesting candidate for an early mediator of thalamocortical interactions because 1) dividing neural progenitor cells and newborn postmitotic neurons already express functional glutamate receptors before the formation of thalamocortical synapses (Haydar et al 2000, Loturco et al 1991, LoTurco et al 1995), and 2) TCAs from sensory thalamic nuclei already show spontaneous electrical activity and calcium spikes even before the axons reach the cortex (Mire et al 2012, Moreno-Juan et al 2017, Uesaka et al 2007). Important roles of glutamate in early brain development are also suggested in human cases where heterozygous mutations of *GRIN1* or *GRIN2B* gene, each of which encodes a subunit of the NMDA receptor, are accompanied by epilepsy, intellectual disabilities and autism spectrum disorders(Fry et al 2018, Lemke et al 2016). Therefore, we examined the early requirement of vesicular glutamate release from TCAs by analyzing mice lacking *vGluT2* in the thalamus.

## Results

### Lack of TCA-derived glutamate causes a downward shift of E14.5-born neurons and alters laminar gene expression in the primary sensory cortex

To investigate the requirement of TCA-derived glutamate in early cortical development, we generated thalamus-specific conditional knockout (cKO) mice for *vGluT2*, a vesicular glutamate transporter. Vesicular glutamate transporters are essential for the packaging glutamate into synaptic vesicles (Tong et al 2007b). In the developing thalamus, *vGluT2* is the only glutamate transporter expressed until E18.5, when *vGluT1* begins to be detected in the principal sensory nuclei (Allen Developmental Mouse Brain Atlas).

We used the *Olig3Cre* driver (Bluske et al 2012, Vue et al 2009, Vue et al 2013) to conditionally delete *vGluT2* in the thalamus. At E16.5, wild-type (WT) brains showed an overlapping expression of vGluT2 protein and the TCA marker NetrinG1 (Fig.1A,A’). In thalamus-specific *vGluT2* cKO mice, vGluT2 expression is lost in NetrinG1-positive TCAs (Fig.1C,C’), confirming efficient early deletion of *vGluT2*.

**Figure 1.**
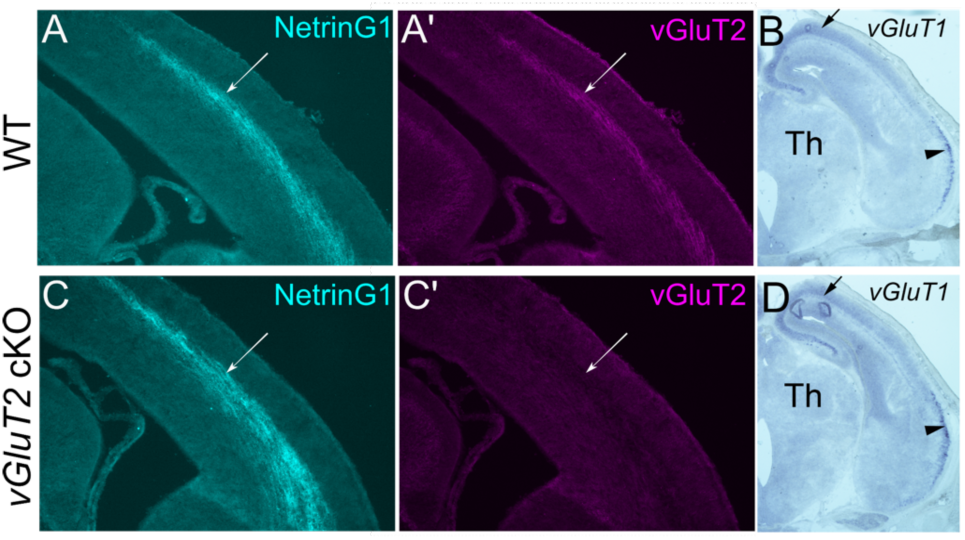
Validation of *vGluT2* cKO mice. A,A’,C,C’. In E16.5 wild-type (WT) cortex, vGluT2 expression overlaps with NetrinG1 in TCA (A, A’). Thalamusspecific *vGluT2* knockout (*vGluT2* cKO) mice show normal TCA projections (C), but TCAs express no vGluT2 protein (C’). **B,D.** At E16.5, *vGluT1*, which encodes another vesicular glutamate transporter, is not detectable in WT thalamus (Th) (B) or *vGluT2* cKO thalamus (D). Medial cortex (arrow) and olfactory cortex (arrowhead) already express *vGluT1* mRNA at this stage.

Another vesicular glutamate transporter, vGluT1, is expressed in principal sensory thalamic nuclei at E18.5 and into postnatal stages (Allen Brain Developing Mouse Brain Atlas). In mice with postnatal deletion of *vGluT2*, glutamatergic TCA synaptic transmission is normal at P9-11 (Li et al 2013), suggesting that *vGluT1* expression in postnatal thalamus compensates for the loss of *vGluT2*. However, at E16.5, we found that *vGluT1* is not yet expressed in the thalamus in either WT or in *vGluT2* cKO mice (Fig.1B,D), indicating that there is no compensatory upregulation of *vGluT1* in the absence of *vGluT2*. Therefore, this model allows us to specifically test whether the absence of vesicular glutamate release during embryogenesis affects neocortical neurogenesis.

Because TCAs establish synaptic connections with layer 4 neurons by approximately P3 (Agmon et al 1995, Kageyama & Robertson 1993), we first analyzed the mutant mice at P1 to assess the role of TCA-derived glutamate prior to synapse formation. At P1, the WT S1 cortex showed segregation of the layer 2/3 marker BRN2 and the layer 4 marker ROR across the future layer boundary (Fig.2A). In contrast, layer 5 contained a subset of CTIP2^+^ cells (35.4±12.3%, n=16) that co-expressed ROR (Fig.2A). In *vGluT2* cKO mice, the number of superficial layer neurons expressing BRN2 and ROR were both reduced compared with WT controls (Fig.2B, grey and yellow). However, the ratio of BRN2^+^ to ROR^+^ cells was increased (Fig.2B, green), indicating a relative shift in cell identity. In layer 5, the number of cells co-expressing ROR and CTIP2 was increased in the cKO cortex. These findings indicate that TCA-derived glutamate is required for the proper generation, specification and/or migration of future layer 4 neurons in the S1 cortex.

**Figure 2.**
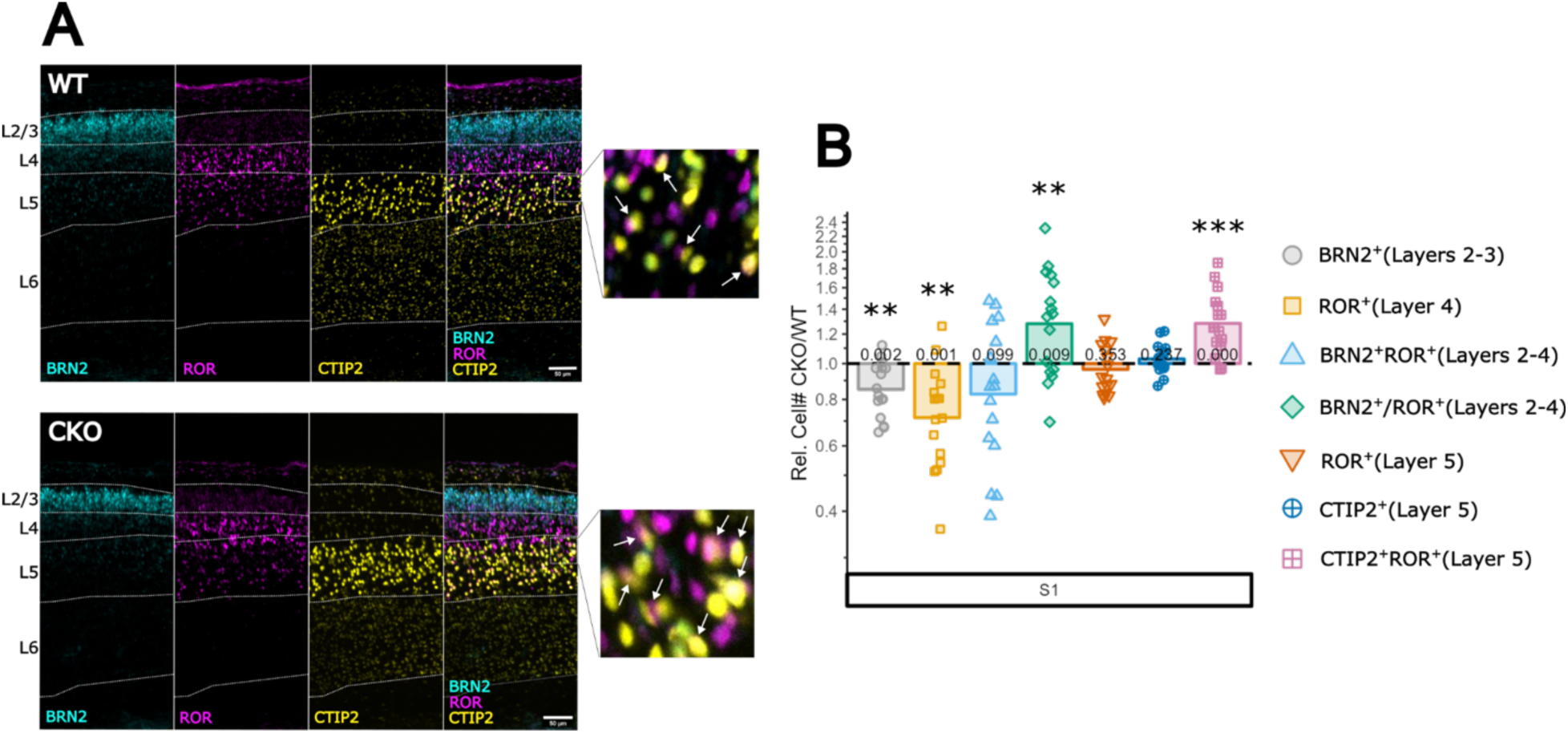
Changes in laminar gene expression in *vGluT2* cKO mice at P1 mice. **A.** Triple staining of littermate WT (top) and vGluT2 cKO (bottom) sections with BRN2 (layers 2 and 3), ROR (layer 4), and CTIP2 (layers 5 and 6). Unlike at P8, many ROR^+^ cells are still found in layer 5, some of which co-express CTIP2. This population largely becomes CTIP2^+^ROR-layer 5 neurons by P8. White arrows highlight cells that co-express ROR and CTIP2 in layer 5. **B.** Analysis of relative cell number for immuno markers (BRN2, ROR and CTIP2) at P1, showing a decrease in BRN2^+^ cells in layer 2/3 as well as ROR^+^ cells in layer 4. The BRN2/ROR ratio is increased, and CTIP2^+^ROR^+^ cells in layer 5 are increased. Data are shown as the relative difference of vGluT2 cKO to WT littermates (dashed line). Points represent individual pairs and bars represent pooled mean. *P < 0.05, **P < 0.01, ***P < 0.001

Because the peak of layer 4 neurogenesis in S1 occurs at E14.5 (Baumann et al 2025), we next examined whether neurons generated at this stage exhibit altered number, radial distribution, or gene expression. To do this, EdU was administered intraperitoneally to pregnant dams at E14.5, and brains were analyzed at P1. We found that the total number of EdU^+^ cells is unchanged in the absence of TCA-derived glutamate (Fig.3B and 3D, grey), indicating that overall cell production at E14.5 is not affected. Notably, in thalamus-specific *Gbx2* knockout mice, which lack most TCAs, the number of EdU^+^ cells labeled at E14.5 is reduced (Monko et al 2022), suggesting while TCAs are required for proper proliferation, glutamate is unlikely to be the key mediator of this specific function. Although the total number of E14.5-born cells was unchanged, the number of EdU^+^ cells located in layer 4, as well as EdU^+^ cells expressing ROR within layer 4, was reduced (Fig.3D, yellow, light blue). In contrast, in layer 5, the total EdU^+^ cells, EdU^+^ROR^+^ cells, and EdU^+^ROR^+^CTIP2^+^ cells were all increased (Fig.3D, green, orange, blue).

**Figure 3.**
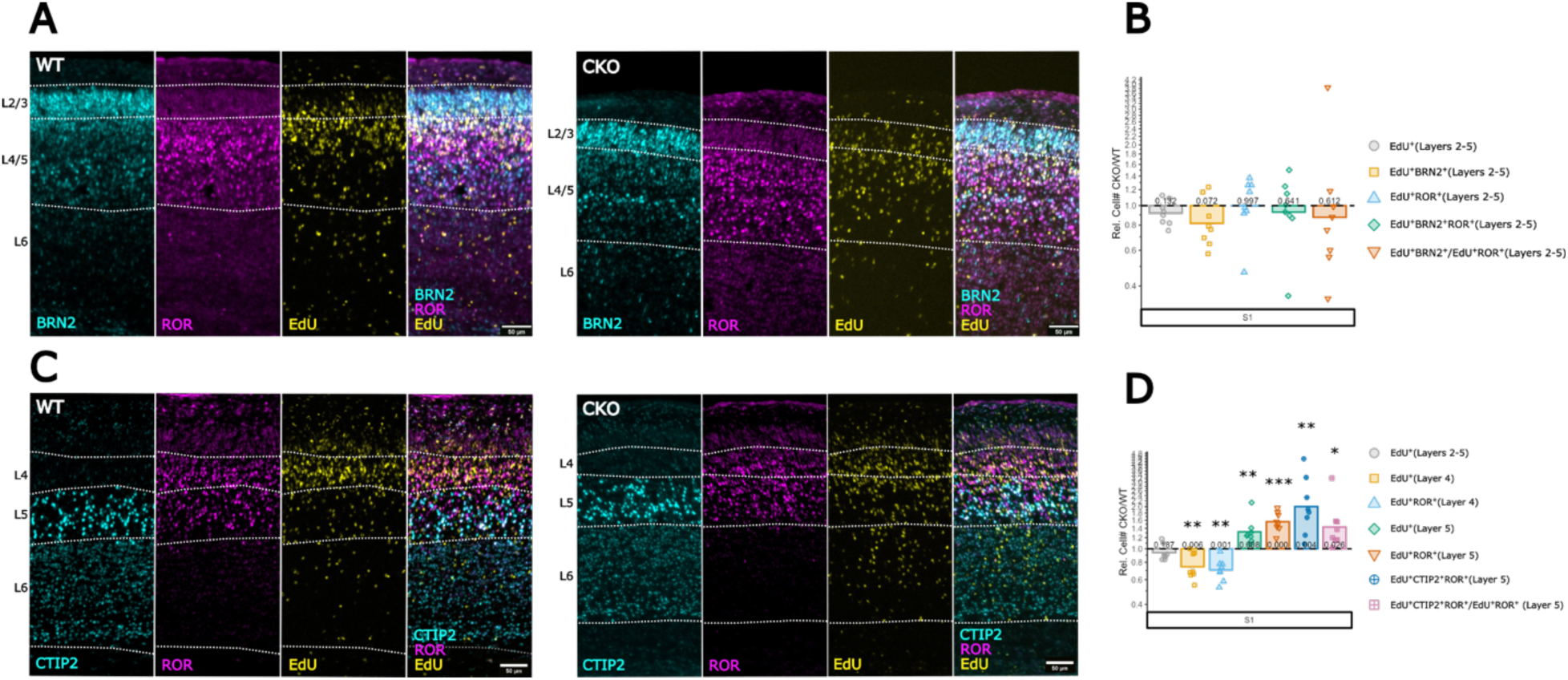
Altered laminar positioning and identity of E14.5-born neurons in *vGluT2* cKO cortex at P1. **A.** Immunostaining of littermate WT (left) and *vGluT2* cKO (right) sections with BRN2 (layers 2 and 3), ROR (layer 4) and EdU (injected at E14.5). **B.** Quantitative analysis of relative cell number for immuno markers (BRN2, ROR, and EdU) at P1, demonstrating no change in EdU^+^BRN2^+^ cells, EdU^+^ROR^+^ cells, and EdU^+^BRN2^+^ROR^+^ cells across all layers (2-5). There was also no change in overall EdU^+^ cells nor the ratio of EdU^+^BRN2^+^/EdU^+^ROR^+^ cells across all layers. **C.** Triple staining of littermate WT (left) and *vGluT2* cKO (right) sections with CTIP2 (layers 5 and 6), ROR (layer 4) and EdU (injected at E14.5). EdU^+^ cells are mainly found in layer 4 in the WT cortex but more EdU^+^ cells are in layer 5 of the cKO cortex. **D.** Analysis of relative cell number for immuno markers (CTIP2, ROR, and EdU) at P1, showing a decrease in EdU^+^ cells and EdU^+^ROR^+^ cells in layer 4. The number of EdU^+^ cells, EdU^+^ROR^+^ cells, and EdU^+^ROR^+^CTIP2^+^ cells were all increased in *vGluT2* cKO at P1. The ratio of EdU^+^ROR^+^CTIP2^+^/EdU^+^ROR^+^ was also increased. The total amount of EdU^+^ cells across all layers (2/3/4/5) was unchanged. For B and D, data are shown as the relative difference of vGluT2 cKO to WT littermates (dashed line). Points represent individual pairs and bars represent pooled mean. *P < 0.05, **P < 0.01, ***P < 0.001.

These results demonstrate that the absence of TCA-derived glutamate leads to a downward shift of E14.5-born neurons by P1. This shift in radial position is accompanied by altered gene expression, including the increased number of neurons in layer 5 that co-express the layer 4 marker ROR and the deep layer marker CTIP2.

We next asked whether TCA-derived glutamate is required for proper radial positioning and gene expression of neurons born before or after E14.5. To address this, we labeled proliferating progenitors at E13.5 or E15.5 using EdU and analyzed the brains at P1.

In WT S1 cortex, the majority of E13.5-born cells were located in layer 5 and 6, overlapping with CTIP2 expression (Fig.4A). In *vGluT2* cKO mice, the total number of E13.5-born cells were unchanged, similar to the E14.5-born population (Fig.4B). Unlike E14.5-born cells, the radial distribution of E13.5-born cells was not altered (Fig.4B), and the number of ROR^+^CTIP2^+^ cells in layer 5 remained unchanged.

**Figure 4.**
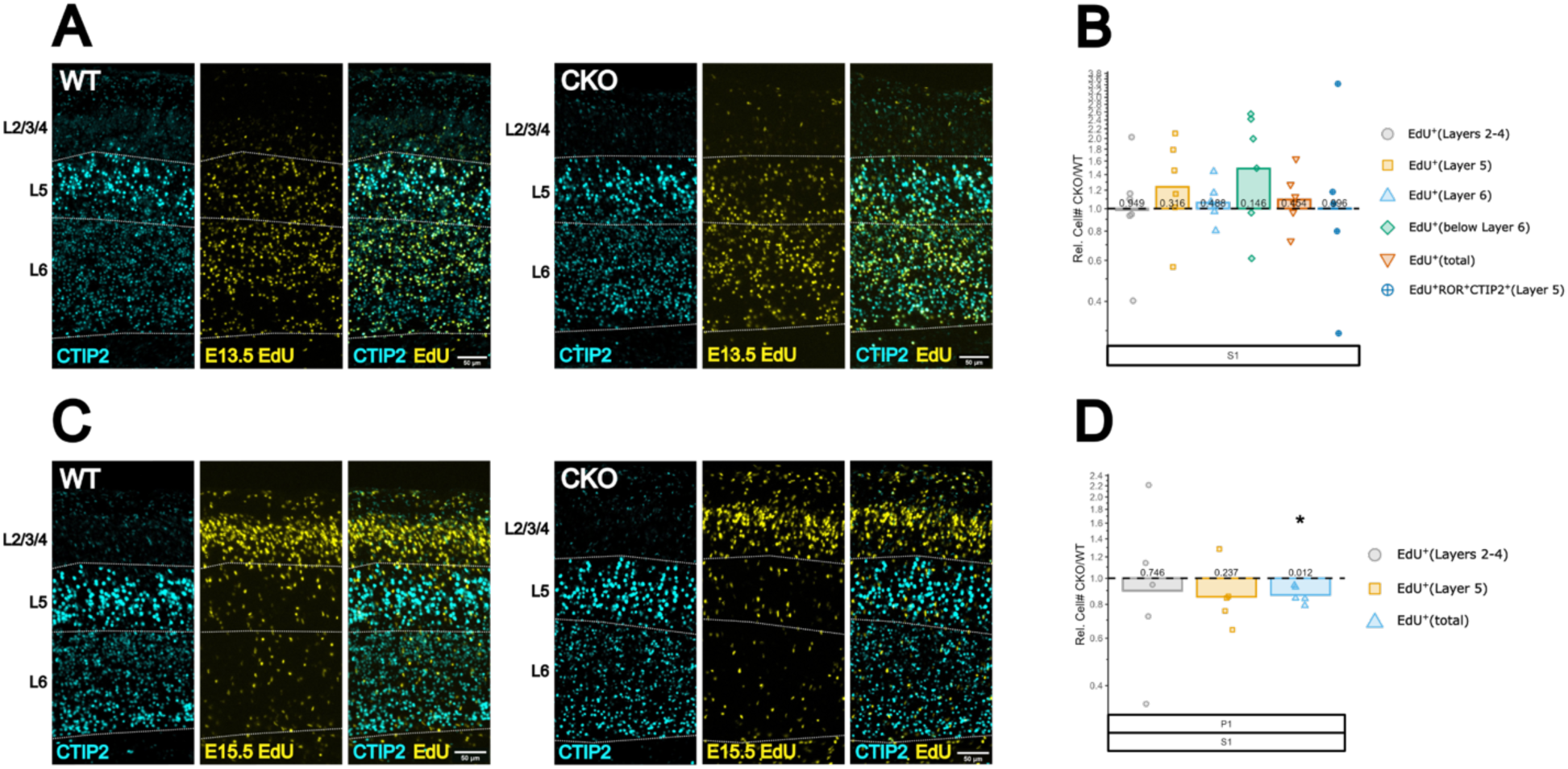
Characterization of the position and identity of E13.5- and E15.5-born cells in *vGluT2* cKO cortex at P1. **A.** Immunostaining of littermate WT (left) and *vGluT2* cKO (right) sections at P1 with CTIP2 (layers 5 and 6) and EdU (injected at E13.5). EdU^+^ cells are mainly found in layers 5 and 6 in WT and vGluT2 cKO cortex. **B.** Analysis of relative cell number for immuno markers (CTIP2 and EdU) at P1, showing no change in total EdU^+^ cells or EdU^+^ cells in layers 5, 6, or below layer 6. There was also no change in the number of EdU^+^ROR^+^CTIP2^+^ cells in layer 5. **C.** Immunostaining of littermate WT (left) and *vGluT2* cKO (right) sections at P1 with CTIP2 (layers 5 and 6) and EdU (injected at E15.5). EdU^+^ cells are mainly found in layers 2, 3, and 4 in WT and vGluT2 cKO cortex. **D.** Analysis of relative cell number for immuno markers (CTIP2 and EdU) at P1, showing no change in the number of EdU^+^ cells present in layers 2-4 or layer 5. The overall number of EdU^+^ cells in all layers (layers 2-5) was decreased in *vGluT2* cKO cortex. For B and D, data are shown as the relative difference of *vGluT2* cKO to WT littermates (dashed line). Points represent individual pairs and bars represent pooled mean. *P < 0.05, **P < 0.01, ***P < 0.001

In WT S1 cortex, most E15.5-born cells were located in layers 2/3 and 4, above the CTIP2-expressing deep layers (Fig.4C). Unlike the E13.5- and E14.5-born cell populations, the total number of E15.5-born cells was reduced in *vGluT2* cKO cortex at P1 (Fig. 4D). This is consistent with an analysis of progenitor cell numbers at E16.5, which showed a reduction in TBR2^+^ intermediate progenitor cells in cKO cortex (Supplemental Fig. 4). However, layer-specific counts of the E15.5-born cohort were unchanged (Fig. 4D), indicating that their radial distribution was not affected.

Together, these results suggest that the requirement of TCA-derived glutamate in regulating the radial positioning and gene expression of excitatory neurons is largely restricted to neurons born at E14.5. Nonetheless, the reduction in the number of E15.5-born cells in the *vGluT2* cKO cortex indicates that TCA-derived glutamate may play a distinct role in regulating the output of late-stage progenitors.

Given the onset of *vGluT1* expression in the principal sensory nuclei of the thalamus at E18.5, we next tested whether the neonatal phonotypes in *vGluT2* cKO mice are still present at later postnatal stages. We found that several phenotypes observed in P1 were no longer present at P4 (Supplemental Fig. 1). For example, the decrease of BRN2^+^ cells in layer 2/3 and the increase in ROR^+^CTIP2^+^ cells in layer 5 were not detected at this stage (Supplemental Fig. 1A,B). In addition, EdU labeling experiments showed that the altered fates of E14.5-labeled neurons observed at P1 were normalized by P4 (Supplemental Fig. 1C,D). Furthermore, barrel field formation was evident in *vGluT2* cKO mice, indicating normal thalamocortical synaptic activity in the postnatal cortex (Supplemental Fig. 1E).

Although a reduction in ROR^+^ cells in layer 4 and an increased ratio of BRN2^+^ to ROR^+^ cells in superficial layers were still observed at P4 (Supplemental Fig. 1A,B), these differences were normalized by P8 (Supplemental Fig.2). Thus, the altered fate of future layer 4 neurons in *vGluT2* cKO mice appears to be transient.

### Mice lacking TCA-derived VGF show distinct fate changes in prospective layer 4 neurons

We previously reported that thalamus-specific *Vgf* cKO mice exhibit a reduction of layer 4 neurons at P8 (Monko et al 2022). Here, we repeated this analysis at P1 using the same parameters applied to *vGluT2* cKO mice. We found that the number of ROR^+^ cells in layer 4 was decreased, whereas the numbers of BRN2^+^ cells and BRN2^+^ROR^+^ cells were increased (Fig.5A).

**Figure 5.**
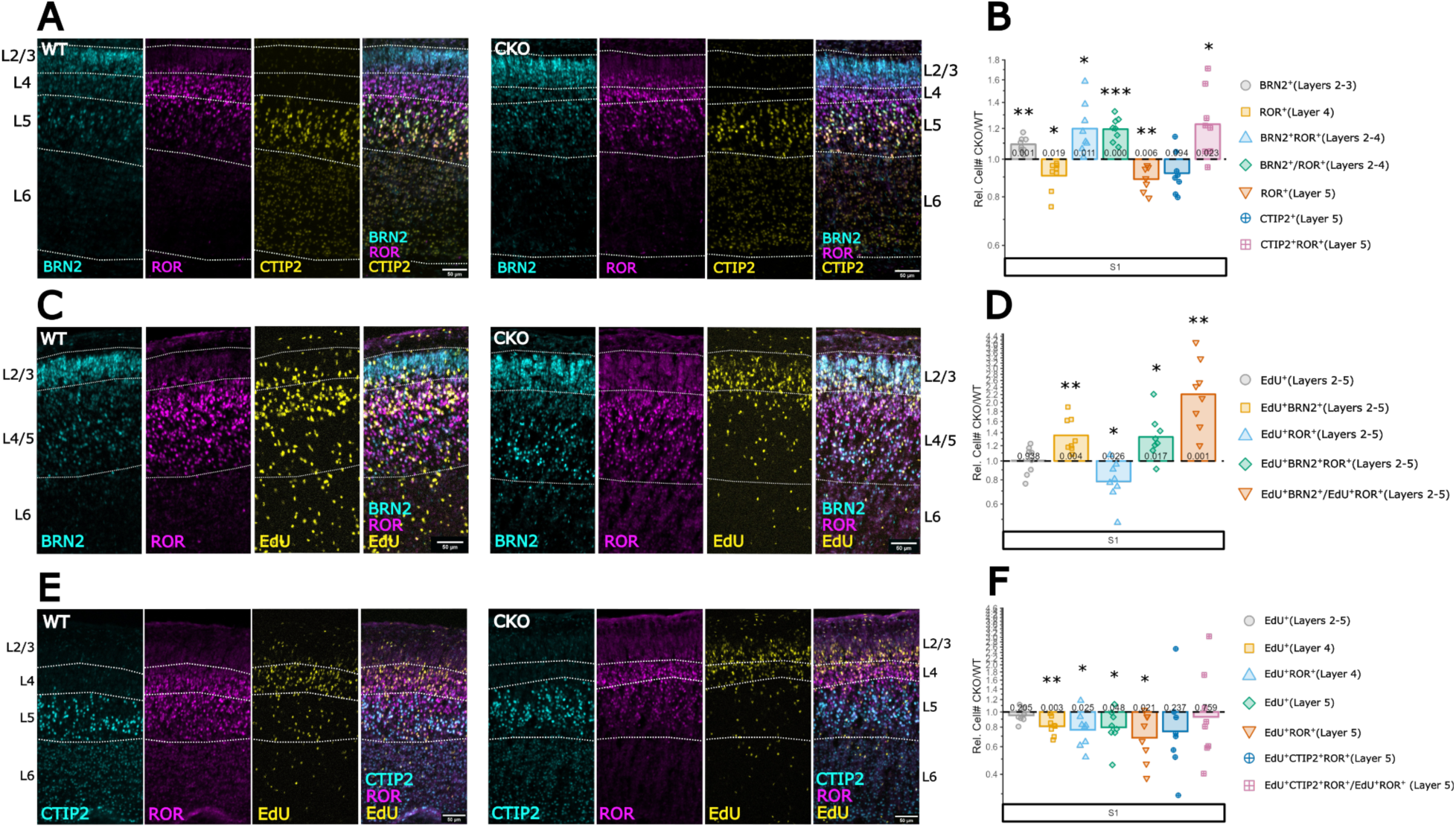
Suppression of TCA-derived VGF shifts E14.5-born cell fate towards layers 2 and 3. **A.** Immunostaining of littermate WT (left) and *Vgf* cKO (right) sections at P1 with BRN2 (layers 2 and 3), ROR (layer 4), and CTIP2 (layers 5 and 6). **B.** Analysis of relative cell number for immuno markers (BRN2, ROR, and CTIP2) at P1, showing an increase in BRN2^+^ and BRN2^+^ROR^+^ cells in layers 2-4. There was a decrease in ROR^+^ cells in both layer 4 and layer 5. The ratio of BRN2^+^/ROR^+^ in layers 2-4 was increased as was the number of ROR^+^CTIP2^+^ cells in layer 5. **C.** Triple staining of littermate WT (left) and *Vgf* cKO (right) sections with BRN2 (layers 2 and 3), ROR (layer 4) and EdU (injected at E14.5). EdU^+^ cells are mainly found in layer 4 in the WT cortex but more EdU^+^ cells are in layers 2 and 3 of the cKO cortex. **D.** Analysis of relative cell number for immuno markers (BRN2, ROR, and EdU) at P1, showing an increase in the number of EdU^+^BRN2^+^ cells and EdU^+^BRN2^+^ROR^+^ cells across all layers (2-5). The number of overall EdU^+^ROR^+^ cells was decreased across all layers and as such the ratio of EdU^+^BRN2^+^/EdU^+^ROR^+^ was increased across all layers. The overall number of EdU^+^cells across all layers was unchanged. **E.** Immunostaining of littermate WT (left) and *Vgf* cKO (right) sections with CTIP2 (layers 5 and 6), ROR (layer 4) and EdU (injected at E14.5). **F.** Analysis of relative cell number for immuno markers (CTIP2, ROR, and EdU) at P1, exhibiting a decrease in EdU^+^ROR^+^ cells in both layer 4 and layer 5. EdU^+^ cells in layers 4 and 5 were also decreased, however, the total amount of EdU^+^ cells across all layers (2/3/4/5) was unchanged. The EdU^+^ROR^+^CTIP2^+^ cell population and ratio of EdU^+^ROR^+^CTIP2^+^/EdU^+^ROR^+^ cells in layer 5 were unchanged in *Vgf* cKO cortex. For B and D and F, data are shown as the relative difference of *Vgf* cKO to WT littermates (dashed line). Points represent individual pairs and bars represent pooled mean. *P < 0.05, **P < 0.01, ***P < 0.001.

In EdU birthdating experiments (labeling at E14.5), the total number of EdU^+^ cells was unchanged, similar to that observed in *vGluT2* cKO mice. However, EdU^+^ cells expressing BRN2 alone or both BRN2 and ROR were increased, whereas EdU^+^ cells expressing ROR alone were decreased. These results indicate that, in the absence of thalamus-derived VGF, E14.5-born neurons undergo a fate shift from ROR^+^ layer 4 cells to BRN2^+^ layer 2/3 cells. This pronounced disruption of neuronal fate in superficial layers was not observed in *vGluT2* cKO mice.

In contrast to *vGluT2* cKO mice, *Vgf* cKO mice showed a decrease in the number of EdU^+^ cells, including those also expressing ROR, in layer 5. There was no significant change in the number EdU^+^ cells co-expressing ROR and CTIP2 in layer 5 (Fig.5E,F). Together, these results demonstrate that the two TCA-derived factors, glutamate and VGF, are required to regulate different aspects of neuronal positioning and gene expression in the embryonic and neonatal sensory cortex.

### Lack of both TCA-derived glutamate and VGF results in additive neocortical phenotypes

The distinct phenotypes observed in *vGluT2* and *Vgf* single cKO mice raise the question of whether these TCA-derived effectors act independently or interactively to influence neocortical neurogenesis. To address this, we examined double conditional knockout (dcKO) mice lacking both *vGluT2* and *VGF*.

Using analytical approaches similar to those applied in the single cKO experiments, we assessed laminar organization and molecular identities of cortical neurons in *vGluT2:Vgf* dcKO mice at P1. For three-way comparisons among WT, *Vgf* single cKO, and *vGluT2:Vgf* dcKO mice from the same litter, we used a linear mixed-effects model (LMM) with Tukey’s correction. In this model, genotype was treated as a fixed effect, and litter was included as a random intercept to account for shared developmental environment and technical variability. The number of ROR^+^ cells in superficial layers was reduced in both *Vgf* single cKO and dcKO cortices compared with WT, with no significant difference between the two mutant groups (Fig. 6B). In dcKO mice, the number of BRN2^+^ neurons in superficial layers was increased relative to WT (Fig. 6C). Notably, statistical significance between single *Vgf* cKO and WT was not detected in the three-way LMM analysis, although direct pairwise comparison between these genotypes revealed a significant difference (Fig. 5). This discrepancy likely reflects the reduced statistical power of multiple-comparison correction and increased variance introduced inclusion of an additional genotype. The number of ROR^+^BRN2^+^ cells in superficial layers was increased in both *Vgf* single cKO and dcKO cortices compared with WT, with no significant difference between the two mutant groups (Fig. 6D). In layer 5, dcKO mice exhibited an increased population of ROR^+^CTIP2^+^ cells compared with both WT and *Vgf* single cKO (Fig. 6E). This increase is consistent with findings in *vGluT2* cKO mice (Fig. 2, Fig. 5B). However, EdU (E14.5)^+^ROR^+^CTIP2^+^ triple-positive cells were increased in layer 5 only in *vGluT2* cKO (Fig. 3D) and not in *Vgf* cKO mice (Fig. 5E). In contrast, the number of ROR^+^ cells in layer 5 was decreased in *Vgf* cKO and dcKO compared with WT (Fig. 5F), whereas there was no change in *vGluT2* cKO.

**Figure 6.**
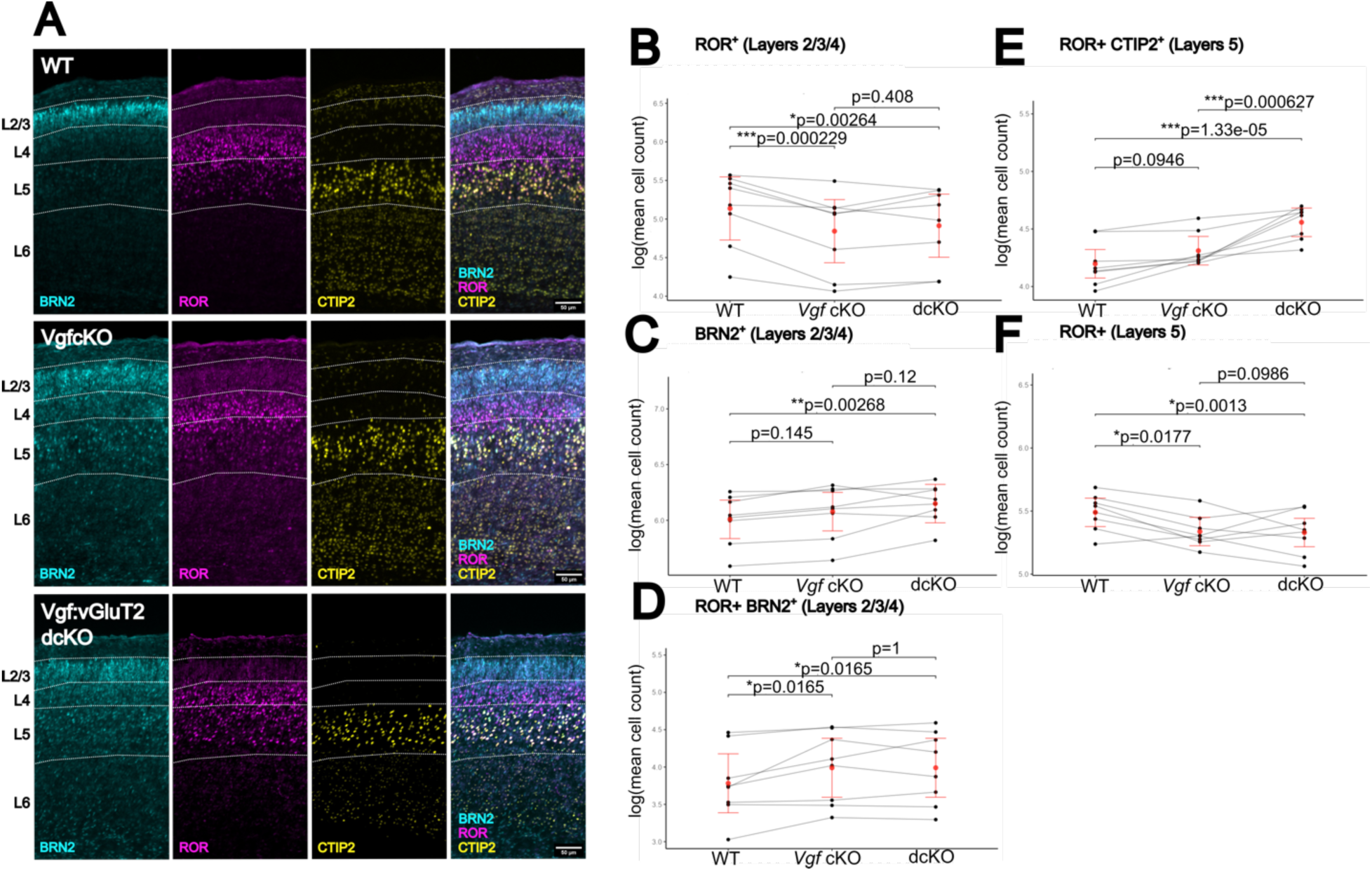
Characterization of *Vgf:vGluT2* dcKO cortex at P1. **A.** Immunohistochemical staining of littermate WT (top), *Vgf* single cKO (middle), and *Vgf:vGluT2* dcKO (bottom) sections at P1 for BRN2 (layers 2 and 3), ROR (layer 4), and CTIP2 (layers 5 and 6). **B-F.** Analysis of cell numbers for immuno markers (BRN2, ROR, and CTIP2) at P1 between the three genotypes using linear mixed-effects model (LMM). LMM analysis was performed using the R package lme4. (B) ROR^+^ cells in layer 4 are significantly decreased in both *Vgf* cKO and dcKO cortex compared with WT, and there is no significant difference between single and double cKO. (C) dcKO cortex shows an increase in BRN2^+^ cells in superficial layers compared with WT. Significance between single cKO and WT is not detected in this analysis, although just comparing between the two still results in significant difference as in Fig.5. This is due to the nature of the three-way comparison using LMM. (D) ROR^+^BRN2^+^ cells in superficial layers are significantly increased in both single cKO and dcKO cortex compared with WT, and there is no significant difference between single cKO and dcKO. (E) ROR^+^CTIP2^+^ cells in layer 5 are unchanged in single *Vgf* cKO but are significantly increased in dcKO compared with WT and single *Vgf* cKO, similar to single *vGluT2* cKO (Fig.4). (F) ROR^+^ cells in layer 5 are decreased in both single cKO and dcKO cortex compared with WT, and there is no significant difference between single cKO and dcKO. Connected lines in each panel represent trios of WT, *Vgf* single cKO and dcKO brains in the same litter. Red points are LMM-estimated marginal means ±95% confidence interval. *P < 0.05, **P < 0.01, ***P < 0.001. Scale bar=50μm.

Together, these findings suggest that TCA-derived glutamate acts independent of VGF to promote the differentiation of ROR^+^CTIP2^+^ neurons, and that the combined loss of glutamate and VGF produces additive effects on neocortical development.

### Mice lacking activity-dependent vesicular release from TCAs show distinct phenotypes from those lacking glutamate or VGF

After synapse formation, neurotransmitter release depends on activity-dependent vesicle fusion mediated by the SNARE complex, and tetanus toxin (TeNT) blocks this process by cleaving SNARE proteins including VAMPs (Baines et al 1999, Yasuda et al 2011, Zhang et al 2008). Release of VGF-derived peptides has been shown to occur via dense core vesicle fusion (Bartolomucci et al 2011), which may also be sensitive to TeNT.

Based on this, we reasoned that if the release of glutamate or VGF from developing TCAs prior to synapse formation depends on TeNT-sensitive vesicular mechanisms, then mis-expression of TeNT in thalamic neurons should phenocopy the effects in *vGluT2* and *Vgf* cKO mice. Consistent with this prediction, mice mis-expressing TeNT under the *Olig3Cre* driver showed a reduction in ROR^+^ layer 4 neurons at P1 (Fig. 7B), similar to both *vGluT2* cKO *and Vgf* cKO mice. The total number of BRN2^+^ cells in superficial layers was unchanged, but the ratio of BRN2^+^ to ROR^+^ cells was increased (Fig. 7B), indicating a shift in neuronal identity.

**Figure 7.**
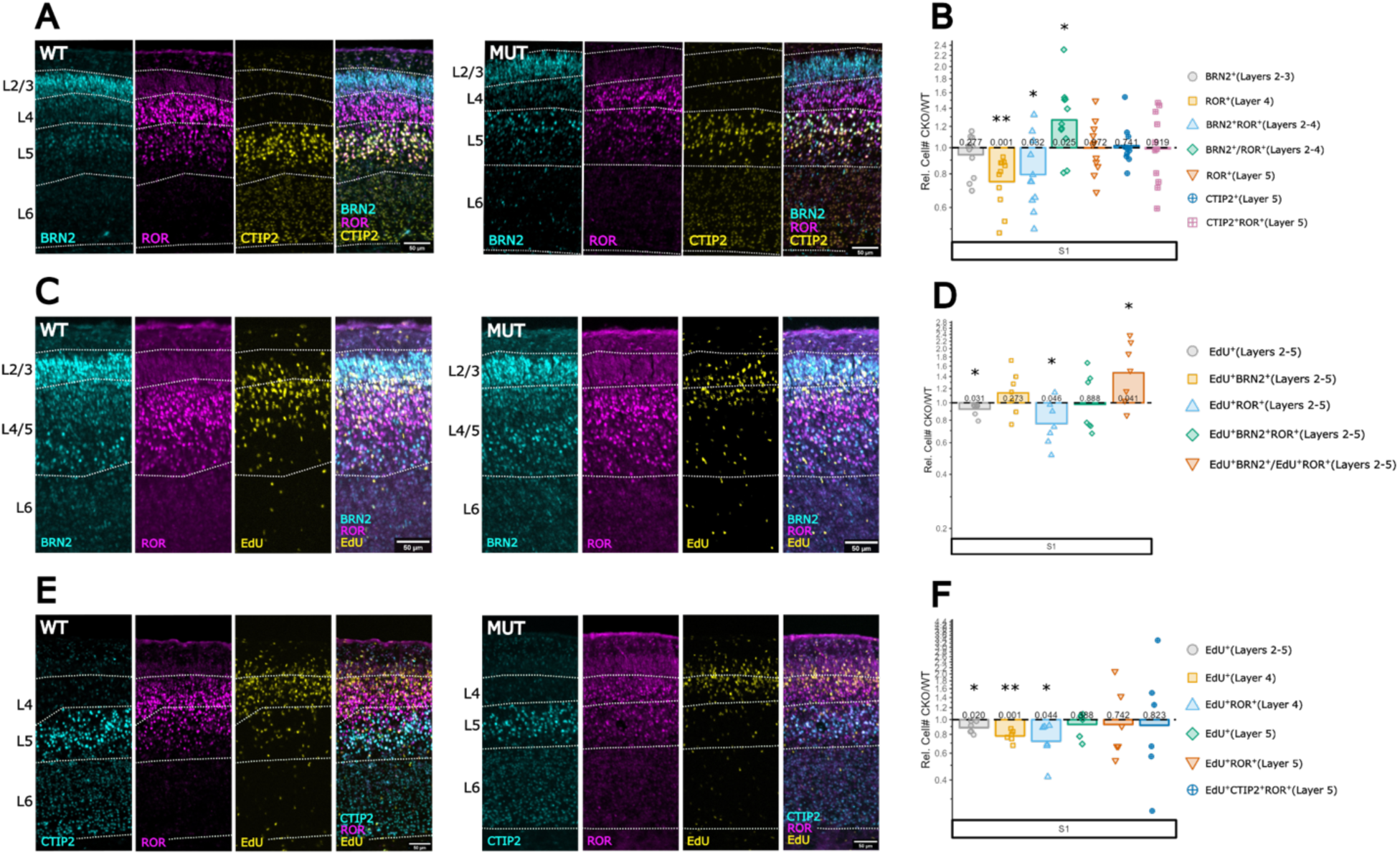
Mice mis-expressing tetanus toxin in TCAs show unique phenotypes from *vGluT2 or Vgf* cKO mice at P1. **A.** Triple staining of littermate WT (left) and TeNT-expressing (right) sections at P1 with BRN2 (layers 2 and 3), ROR (layer 4), and CTIP2 (layers 5 and 6). **B.** Analysis of relative cell number for immuno markers (BRN2, ROR, and CTIP2) at P1, showing a decrease in ROR^+^ and BRN2^+^ROR^+^ cells in superficial layers. The number of BRN2^+^ cells in superficial layers was not changed, however, the ratio of BRN2^+^/ROR^+^ cells in layer 2-4 was increased. **C.** Immunostaining of littermate WT (left) and TeNT-expressing (right) sections with BRN2 (layers 2 and 3), ROR (layer 4) and EdU (injected at E14.5). **D.** Analysis of relative cell number for immuno markers (BRN2, ROR, and EdU) at P1, exhibiting a decrease in EdU^+^ROR^+^ cells in layers 2-5 and an increase in the ratio of EdU^+^BRN2^+^/EdU^+^ROR^+^ cells in all layers. The number of EdU^+^BRN2^+^ cells in all layers was unchanged and the total number of EdU^+^ cells in all layers was decreased. **E.** Immunostaining of littermate WT (left) and TeNT-expressing (right) sections with CTIP2 (layers 5 and 6), ROR (layer 4) and EdU (injected at E14.5). **F.** Analysis of relative cell number for immuno markers (CTIP2, ROR, and EdU) at P1, showing a decrease in EdU^+^ cells and EdU^+^ROR^+^ cells in layer 4. There was no change in layer 5 cell populations. The total number of EdU^+^ cells in all layers was decreased. For B, D, and F, data are shown as the relative difference of TeNT-expressing to WT littermates (dashed line). Points represent individual pairs and bars represent pooled mean. *P < 0.05, **P < 0.01, ***P < 0.001.

Unlike *vGluT2* cKO, P1 TeNT-expressing mice showed no change in the number of ROR^+^ cells or CTIP2^+^ROR^+^ cells in layer 5 at P1 (Fig. 7B). In EdU birthdating experiments with labeling at E14.5, TeNT-expressing mice exhibited a reduction in total EdU^+^ cells (Fig. 7D), a phenotype not observed in either *vGluT2* cKO (Fig. 3A,B) or *Vgf* cKO (Fig. 5E,F) mice. TeNT-expressing mice also had displayed reduced numbers of EdU^+^ cells and EdU^+^ROR^+^ cells in layer 4, while EdU^+^ cell populations of layer 5 remained unchanged (Fig. 7F). There was no change in EdU^+^BRN2^+^ cells in superficial layers; however, the ratio of EdU^+^BRN2^+^ to EdU^+^ROR^+^ was increased (Fig. 7D), suggesting a shift in fate between BRN2^+^ and ROR^+^ cells.

Taken together, the distinct phenotypes observed in TeNT-expressing mice compared with *vGluT2* cKO and *Vgf* cKO mice indicate that early release of glutamate or VGF from TCAs does not rely on TeNT-sensitive vesicular mechanisms. A previous *in vitro* study showed that glutamate release from growing axons of embryonic hippocampal neurons before synapse formation is not disrupted by TeNT (Verderio et al 1999), which is consistent with our finding. Furthermore, the reduction in overall EdU^+^ cells in TeNT-expressing cortex suggests that progenitor proliferation depends on other TeNT-sensitive processes independent of TCA-derived glutamate or VGF.

## Discussion

In this study, we found that mice lacking vGluT2 specifically in the thalamus exhibit altered positioning and gene expression of prospective layer 4 neurons in neonatal somatosensory cortex. The downward shift of E14.5-born neurons from layer 4 to layer 5 was already evident prior to establishment of direct synaptic input from TCAs. In addition to this change in radial position, there was an increase in the number of E14.5-born neurons in layer 5 that co-express ROR and CTIP2, as well as an increased number of EdU^+^ROR^+^ cells that also express CTIP2 in the cKO cortex. Because ROR and CTIP2 are largely segregated to layer 4 and layer 5, respectively, in the wild-type postnatal cortex, their increased co-expression in *vGluT2* cKO cortex suggests a critical role of TCA-derived glutamate in regulating gene expression and cell migration required for proper layer 4 fate specification.

CTIP2 is a key regulator of subcortically projecting neurons in layer 5 (Arlotta et al 2005, Chen et al 2005), and its expression is directly suppressed by SATB2, which is expressed in intracortically projecting neurons across layers 2 to 5 (Alcamo et al 2008). Therefore, it is possible that the increased co-expression of ROR and CTIP2 in *vGluT2* cKO cortex is due to reduced expression or activity of SATB2. However, we find that although ROR^+^SATB2^+^ double-positive cells were decreased in superficial layers, SATB2^+^CTIP2^+^ cells in layers 5 and 6 were unchanged (Supplemental Fig. 3). Thus, the increase in ROR^+^CTIP2^+^ cells in deep layers is unlikely to result from altered SATB2 function.

The altered fate of prospective layer 4 neurons may result from the direct action of glutamate on newborn postmitotic neurons. Tracking with EdU labeling indicates that E14.5-born neurons migrate past the TCA layer between E16.5 and E18.5 (Monko et al 2022), allowing possible interactions between TCAs and the immature migrating neurons. Alternatively, TCA-derived glutamate may influence the temporal state of radial glia (Telley et al 2019, Telley et al 2016, Vitali et al 2018), whose apical processes extend through the region traversed by incoming TCAs, allowing potential interactions.

Notably, some glutamate receptors are predominantly expressed in postmitotic cortical neurons (e.g., *Grin1*, *Grin2b*, *Grm5*), whereas others are expressed both in neurons and progenitor cells (Grik2, Grik5, Gria2)(Allen Developing Mouse Brain Altas), suggesting multiple potential cellular targets for glutamate signaling.

An intriguing aspect of our findings is that neuronal fate changes were specific to the E14.5-born cohort; neurons labeled by EdU at E13.5 or E15.5 displayed normal radial positioning and gene expression in the cKO cortex (Fig.4). This stage-specific requirement suggests that the spatiotemporal relationship between TCAs and either newborn neurons or progenitor cells is critical for afferent regulation of neocortical neurogenesis. The reduction of cell production observed in the E15.5-born cohort in the absence of TCA-derived glutamate (Fig. 4C,D) suggests that glutamate may also regulate progenitor proliferation at later stages, independently of its role in fate specification. Further analysis, including transcriptomic profiling and assessment of radial glial temporal states (Telley et al 2018, Vitali et al 2018) will be necessary to identify the specific cell populations affected by TCA-derived glutamate.

By P4, some of the neuronal fate phenotypes in *vGluT2* cKO cortex at P1 began to resolve, including the increase in ROR^+^CTIP2^+^ double-positive cells in layer 5. Barrel formation was also evident at this stage, suggesting that glutamatergic signaling from TCAs is restored in postnatal *vGluT2* cKO mice. By P8, the major neuronal phenotypes observed at P1 were no longer detectable. These findings are consistent with previous work comparing thalamus-specific *vGluT2* cKO mice and mice that lack both *vGluT1* and *vGluT2* in the postnatal cortex (Li et al 2013). Because vGluT1 expression begins in the thalamus after E18.5, the restoration of glutamate release from TCAs in the postnatal period likely underlies the recovery of layer 4 phenotypes. It will be important to determine whether these transient developmental defects have lasting functional consequences in later life, as similar “sleeper effects” have been reported in both neurodevelopmental and neurodegenerative disorders (Braz et al 2022, Magno et al 2021).

A comparison between mice lacking TCA-derived glutamate and those lacking TCA-derived VGF reveals strikingly distinct phenotypes. In *vGluT2* cKO mice, E14.5-born cells were shifted downward into layer 5 and exhibited increased co-expression of ROR and CTIP2. In contrast, in *Vgf* cKO mice, E14.5-born neurons were shifted upward into layer 2/3, with an increase in EdU^+^BRN2^+^ cells and a decrease in EdU^+^ and EdU^+^ROR^+^ cells in layer 5 (Fig.5). These contrasting phenotypes indicate that glutamate and VGF regulate distinct aspects of neuronal specification during early neocortical development, although both are required for generating the proper number of layer 4 neurons in sensory cortex.

It remains unclear whether TCA-derived glutamate and VGF act independently or interactively. In double conditional mutant mice lacking both vGluT2 and VGF, all phenotypes observed in *Vgf* cKO cortex were retained (Fig. 6), including features not present in *vGluT2* single cKO mice, such as an increased BRN2^+^ cells in layer 2/3 and decreased ROR^+^ cells in layer 5 (Fig. 2B). Moreover, a three-way comparison between *Vgf* single cKO, dcKO and WT mice revealed that the increase in ROR^+^CTIP2^+^ double-positive cells in layer 5 was further enhanced in dcKO compared with *Vgf* single cKO mice (Fig. 6E), indicating an additive effect of *vGluT2* deletion.

In other systems, such as adult hippocampal progenitors (Thakker-Varia et al 2014) and spinal neurons (Skorput et al 2018), the VGF-derived peptide TLQP-62 enhances neural progenitor cell proliferation and Ca^2+^ responses, respectively, both in a manner dependent on glutamate signaling. In contrast, our results reveal distinct and largely non-overlapping phenotypes in mice lacking TCA-derived glutamate or VGF, with additive effects observed in double mutants. These findings suggest that, in embryonic and neonatal sensory cortex, TCA-derived glutamate and VGF TCAs act largely independently, likely targeting in different cellular populations and/or signaling pathways in a sequential or parallel manner.

Proper early fate specification of excitatory is disrupted in cortical organoid models from individuals with *DISC1* mutations (Qian et al 2020), which area associated with schizophrenia, major depression, and autism (Brandon & Sawa 2011). In addition, thalamocortical assembloid models of *CACNA1G* mutation and 22q11.2 microdeletion, both of which are associated with neurodevelopmental disorders, exhibit abnormal thalamocortical connectivity (Kim et al 2024, Leyva-Diaz et al 2024, Shin et al 2024). Thus, elucidating how major neocortical neuronal types are specified and how thalamocortical axons influence this process is essential for understanding the mechanisms of neurodevelopmental disorders and for developing therapeutic strategies.

## Acknowledgments

We thank Stephen Salton and Martyn Goulding for *Vgf* floxed mice and TeNT-expressing mice, respectively. Reetu Gurung provided assistance in genotyping mice. This work was supported by funds from NINDS (1R21NS130271, 1R21NS117978), Wallin Neuroscience Discovery grant, and University of Minnesota.

## Materials and Methods

### Mice

Mouse husbandry and experimentation were done in accordance with the Institutional Animal Care and Use Committee at the University of Minnesota. Noon on the day in which a vaginal plug was found was designated as embryonic day 0.5 (E0.5). The day of birth (E19.5) was defined as postnatal day 0 (P0). To generate mice with a conditional deletion of *vGluT2* in thalamocortical axons (TCAs), *Olig3^Cre/+^*; *vGluT2^flox/+^* mice (Tong et al 2007a, Vue et al 2009) were bred with *Olig3^+/+^*; *vGluT2^flox/flox^* mice. *Olig3^+/+^*; *vGluT2^flox/(flox^ ^or^ ^+)^* pups were used as wildtype (WT) controls and were compared to *Olig3^Cre/+^*; *vGluT2^flox/flox^* conditional knockout (cKO) littermates. *vGluT2^flox/flox^* mice were obtained from Jackson Laboratory (Strain #012898) and were kept in the C57BL/6 background. For mice with a conditional deletion of *Vgf* in TCAs, *Olig3^Cre/+^*; *Vgf^flox/(flox^ ^or^ ^+)^* mice were bred with *Olig3^+/+^*; *Vgf^flox/flox^* mice. *Olig3^+/+^*; *Vgf^flox/(flox^ ^or^ ^+)^* mice were used as WT controls for comparison with *Olig3^Cre/+^*; *Vgf^flox/flox^* cKO littermates. *Vgf^flox/flox^* mice were received from Dr. Stephen Salton (Lin et al 2015) and were kept in the C57BL/6 background. To generate mice with a combined deletion of both *vGluT2 and Vgf* in TCAs, *Olig3^Cre/+^*; *Vgf^flox/+^*; *vGluT2^flox/+^* mice were bred with *= _Vgf_^flox/flox^*_; *vGluT2*_*^flox/+^* _mice. *Olig3*_*^+/+^*_; *Vgf*_*^flox/(flox or +)^*_; *vGluT2*_*^flox/(flox or +)^* mice were used as WT control and were compared to *Olig3^Cre/+^*; *Vgf^flox/flox^*; *vGluT2^flox/+^* single *Vgf* cKO and *Olig3^Cre/+^*; *Vgf^flox/flox^*; *vGluT2^flox/flox^* double cKO (dcKO) littermates. Thalamus-specific mis-expression of tetanus toxin light chain (TeNT) was achieved by breeding *Olig3^Cre/+^*; *Rosa26^stop-+/+^* mice with *Olig3^+/+^*; *Rosa26^stop-TeNT/TeNT^*mice (Lanuza et al 2004, Larsen et al 2019). *Olig3^+/+^*; *Rosa26^stop-TeNT/+^*mice were used at WT control for comparison with *Olig3^Cre/+^*; *Rosa26^stop-TeNT/+^* TeNT mis-expressing littermates. *Rosa26^stop-TeNT/+^*were obtained from Dr. Martyn Goulding (Zhang et al 2008) and were kept in the C57BL/6 background. For all conditional knockout experiments, mice were matched with littermates of WT and CKO to control for developmental differences between litters. Litters without both a WT and CKO pup were not used for analysis. Both male and female pups were used for all analyses.

### EdU

For experiments involving 5-ethynyl-2’-deoxyuridine (EdU; Carbosynth, Compton, UK), mice were intraperitoneally injected with 50µg EdU/g bodyweight in a saline solution. EdU injections were done between 9am and 1pm at the desired stage. For cell fate analysis, pregnant dams were injected with EdU at E13.5, E14.5, or E15.5 and sacrificed at the desired stage (P1, P4, or P8).

### Tissue preparation

Embryos (E16.5) were intracardiacally perfused with 4% paraformaldehyde (PFA), followed by brain extraction and post-fixation in PFA for 30 minutes before immersion in a 30% sucrose/0.1M phosphate buffer (PB) solution overnight at 4°C. Brains were then frozen in TissueTek OCT compound on dry ice and stored at -80°C until use. Postnatal (P1, P4, and P8) pups were processed in the same way except that the extracted brains were post-fixed for 45 minutes (P1) or 60 minutes (P4 and P8). Prior to PFA perfusion, P8 brains were first perfused with 0.1M PB. To obtain tangential sections through layer 4 (Supplemental Fig. 1E), cortices were extracted after perfusion fixation and sandwiched between two slides with a 1mm-thick spacer for 2-3 hours before immersion in 30% sucrose.

### Immunohistochemistry and EdU detection

Immunohistochemistry protocols were based on (Monko et al 2022). Brains were sectioned with a cryostat at a thickness of 20µm (E16.5, P1), 30µm (P4), or 40µm (P8). Paired littermate WT and CKO brains were sectioned onto the same slides to control for the variability of immunohistochemical staining between slides. Following cryosectioning, sections on slides were dried on a slide warmer for 30 minutes. Slides were then washed with 0.1M phosphate-buffered saline (PBS) for 5 minutes and blocked in 3% donkey serum/0.3% Triton X-100/PBS solution for 1 hour at room temperature. Slides stained for transcription factors were immersed in a boiled 10mM citrate buffer (pH 6) for 5 minutes and washed with PBS twice for 3 minutes prior to blocking. Slides were then treated with primary antibodies at the appropriate dilutions and incubated overnight at 4°C. The following primary antibodies were used:

| Protein | Dilution | Boiled<br>? | Manufacturer | Catalog # | Species |
| --- | --- | --- | --- | --- | --- |
| BRN2 | 200 (100 w/EdU) | Yes | Thermo Scientific | PA5-30124 | Rabbit |
| ROR | 100 (50 w/ EdU) | Yes | R&D Systems | PPH392500 | Mouse |
| CTIP2 | 200 | Yes | Abcam | ab18465 | Rat |
| vGluT2 | 200 | No | Synaptic Systems | 135403 | Rabbit |
| NetrinG1 | 100 | No | R&D Systems | AF1166 | Goat |
| VAMP2 | 100 | No | Synaptic Systems | 104 202 | Rabbit |
| VGF(R15) | 50 | No | Santa Cruz | sc-10383 | Goat |
| PAX6 | 100 | Yes | BD Biosciences | 561462 | Mouse |
| TBR2 | 200 | Yes | Abcam | ab23345 | Rabbit |
| SATB2 | 200 | Yes | Abcam | ab92446 | Rabbit |

Following overnight primary antibody incubation, slides were rinsed three times with PBS for 10 minutes each and incubated for 1 hour at room temperature with fluorochrome-conjugated secondary antibodies (Cy2, Cy3, or Cy5, from Jackson ImmunoResearch). After two 10-minute washes in PBS, slides were counterstained with DAPI, dehydrated in ascending concentrations of ethanol, cleared in xylene, and mounted in DPX mounting medium (Sigma-Aldrich).

For visualization of EdU, following secondary antibody treatment and PBS washes, slides were treated with a solution of 100mM Tris (pH 7.5)/4mM CuSO_4_/100mM sodium ascorbate (freshly prepared)/5µM sulfo-cy5 azide (Lumiprobe) for 20 minutes at room temperature. Slides were then rinsed with PBS three times for 5 minutes each and counterstained, dehydrated, and mounted per the immunohistochemistry protocol above.

### Imaging

Slides that went through immunohistochemical protocols were then imaged using an E800 upright microscope (Nikon) and an Infinity8-9M camera (Teledyne Lumenera). Images were taken using the Infinity Analyze 7 software (Teledyne Lumenera) as 12-bit grayscale TIFF files. Exposure times were selected to maximize the signal-to-noise ratio and were kept identical for slides with the same antibody and within the same matched pair. To avoid cross-channel excitation, slides treated with multiple antibodies were imaged in the order of Cy3, Cy2, and Cy5.

### Binning

Images were processed within the FIJI distribution of ImageJ (Schindelin et al 2012). Individual TIFF files for each channel were opened and merged into a composite stack. A custom FIJI macro was used to overlay a rectangular region of interest (ROI) over each composite stack from pial surface to lateral ventricle (https://github.com/davenrock01/ThalamocorticalGlutamatePaper/). ROIs were overlaid at a width of 200µm for visual and motor cortex sections and 400µm for somatosensory cortex sections.

To ensure a consistency of bin positions within each region, an embryonic and early postnatal mouse atlas (Paxinos et al 2006) was used to determine criteria for bin positioning. We considered the S1 (primary somatosensory cortex)-containing section to be the caudal-most section where the full hippocampus spanned horizontally. The lateral border of the lateral ventricle was used as the lateral border of the S1 bin.

### Thresholding

Binned images were thresholded using a custom FIJI macro (https://github.com/davenrock01/ThalamocorticalGlutamatePaper/). This custom macro batch-processed all images within a matched littermate pair with identical thresholding parameters. This thresholding process removes bias and inconsistency associated with manual cell counts. The specifics of this thresholding macro are outlined in (Monko et al 2022) .

### Colocalization analysis

To detect individual cells that express multiple markers, a custom FIJI macro was used (https://github.com/davenrock01/ThalamocorticalGlutamatePaper/). This custom macro identified cells that were double- or triple-positive for different antibodies with a minimum overlap percentage of 35%. The specifics of this colocalization macro are outlined in (Monko et al 2022).

### Cell counting

Thresholded images were counted using the “Analyze Particles” function in FIJI. Cell counts were restricted to cells within the designated bins. For layer-specific cell counts, layer boundaries were drawn manually in FIJI. Cell counts were stored in comma-separated values (.csv) files. Ratios of cell counts (e.g., X^+^Y^+^ / Y^+^) were calculated on a per-section basis.

### Experimental Design and Statistical Analysis

For comparing cKO and WT mice, littermates were matched for experiments and statistical analysis to control for a small variation of developmental timing and variability in immunohistochemistry. Both male and female pups were used for analysis. For each mouse and neocortical region, at least three sections were imaged and analyzed. Data from the same neocortical region within each mouse was averaged as a representative cell count for the region. At least five matched pairs were used for each analysis. A matched ratio t-test was used to compare the logarithm of the average representative cell counts between single cKO and WT littermates. A custom R code was used to perform these analyses and generate plots from cell count datasets (https://github.com/davenrock01/ThalamocorticalGlutamatePaper/). For comparing *Vgf:vGluT2* double cKO (dcKO), *Vgf* single cKO, and wildtype (WT) controls within the same litter, we used linear mixed-effect model (LMM). Genotype was modeled as a fixed effect, while litter was included as random intercepts to account for non-independence due to shared developmental environment and technical variance. LMM analysis was performed using the R package lme4.

### Code and Macros

All custom FIJI macros and R codes are available on a Github repository: https://github.com/davenrock01/ThalamocorticalGlutamatePaper/

---

**Supplemental Figure 1.**
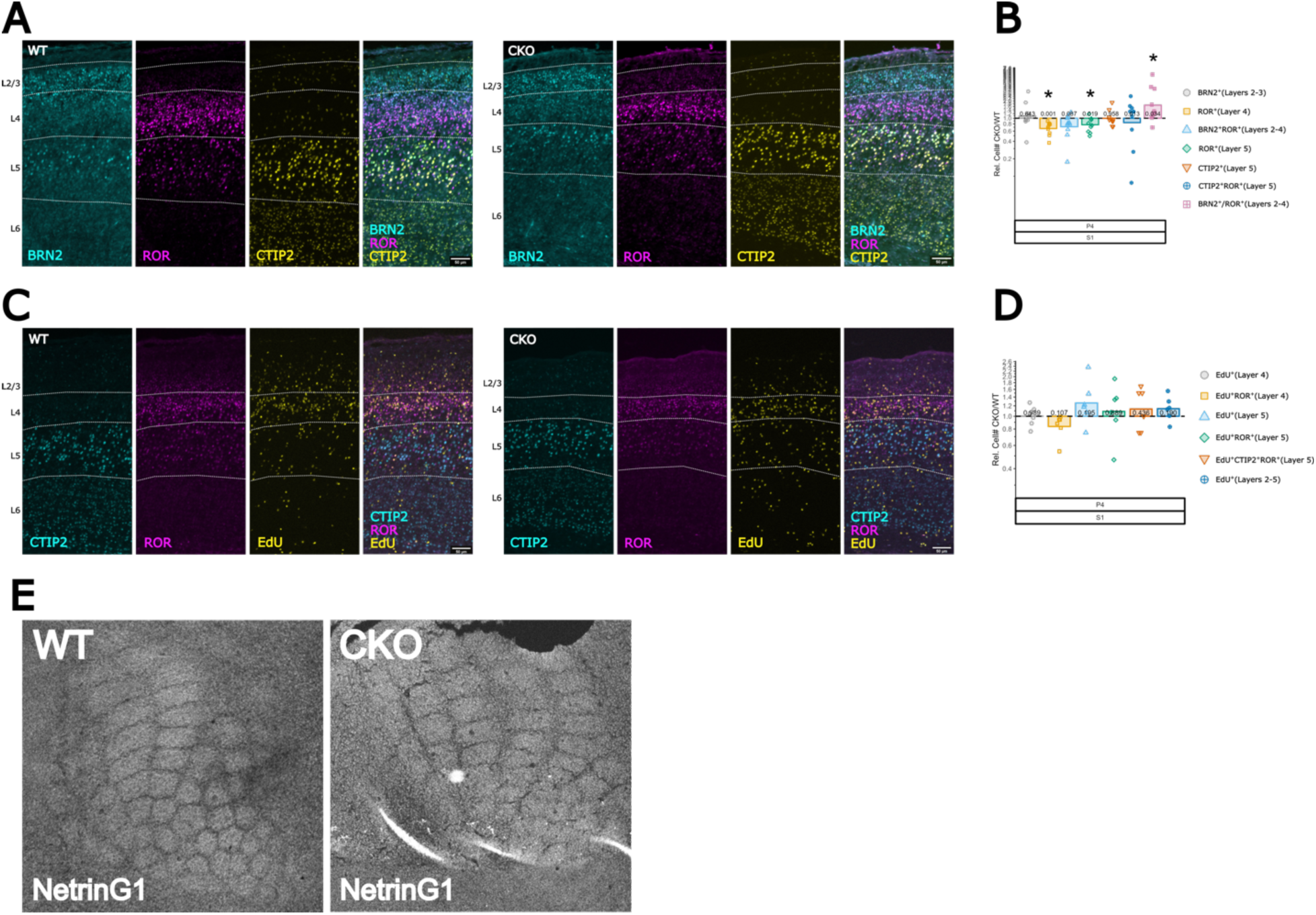
Partial recovery of neuronal fate phenotypes of P1 *vGluT2* cKO mice at P4. **A, B.** Triple staining of littermate WT (left) and *vGluT2* cKO (right) sections with BRN2 (layers 2 and 3), ROR (layer 4), and CTIP2 (layers 5 and 6) at P4. Reduction of ROR^+^ cells in layer 4 is still observed at P4, so is the increased BRN2^+^ to ROR^+^ ratio in superficial layers. However, increase in CTIP2^+^ROR^+^ cells in layer 5, which is seen at P1, is no longer present at P4. **C, D.** EdU labeling experiments show that the fate of E14.5-born cells was also restored by P4. **E.** Immunostaining of tangential sections through layer 4 with anti-NetrinG1 antibody shows that by P4, vGluT2 cKO cortex has already developed barrels in S1, similar to WT controls. *P < 0.05, **P < 0.01, ***P < 0.001.

**Supplemental Figure 2.**
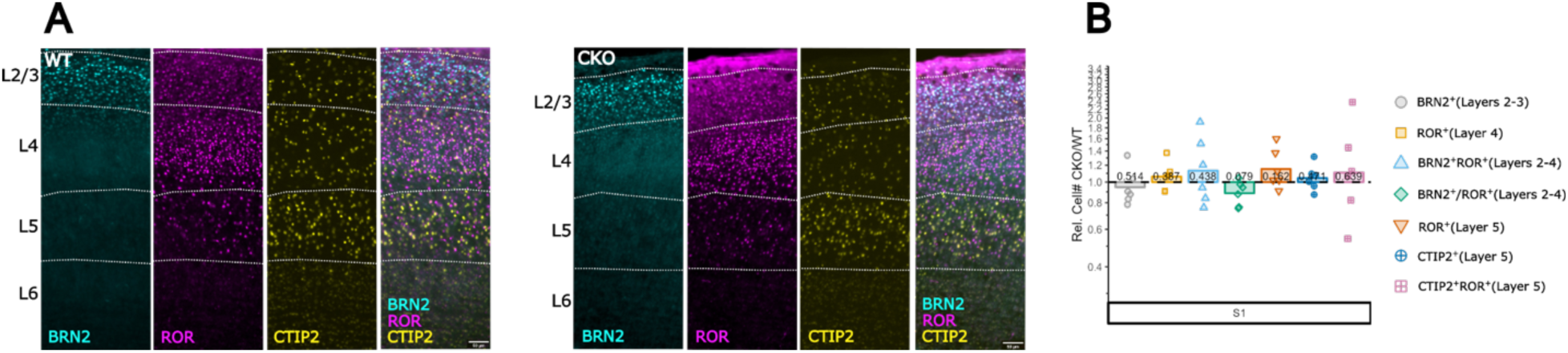
Recovery of neuronal fate phenotypes of P1 *vGluT2* cKO mice at P8. **A.** Triple staining of littermate WT (left) and *vGluT2* cKO (right) sections with BRN2 (layers 2 and 3), ROR (layer 4), and CTIP2 (layers 5 and 6) at P8. At this stage, there are few cells that co-express ROR and CTIP2. However, a small population of ROR^+^CTIP2-cells resides in layer 5. **B.** Analysis of relative cell number for immuno markers (BRN2, ROR and CTIP2) at P8, showing no change in BRN2^+^ cells, ROR^+^ cells, CTIP2^+^ cells, and double-positive cell populations. Data are shown as the relative difference of *vGluT2* cKO to WT littermates (dashed line). Points represent individual pairs and bars represent pooled mean. *P < 0.05, **P < 0.01, ***P < 0.001.

**Supplemental Figure 3.**
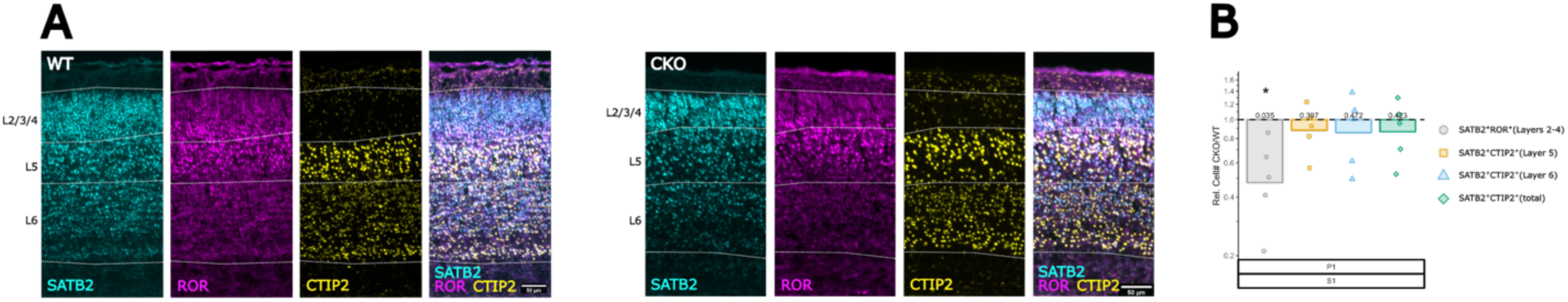
Lack of thalamus-derived glutamate does not alter SATB2 co-expression in cortical neurons at P1. **A.** Triple staining of littermate WT (left) and vGluT2 cKO (right) sections with SATB2 (layers 2-4), ROR (layer 4), and CTIP2 (layers 5 and 6) at P1. At this stage, there are many cells that co-express SATB2 and ROR as well as SATB2 and CTIP2. **B.** Analysis of relative cell number for immuno markers (SATB2, ROR and CTIP2) at P1, showing a decrease in SATB2^+^ROR^+^ cells in layers 2-4 and no change in total or layer-specific SATB2^+^CTIP2^+^ cell populations. Data are shown as the relative difference of *vGluT2* cKO to WT littermates (dashed line). Points represent individual pairs and bars represent pooled mean. *P < 0.05, **P < 0.01, ***P < 0.001.

**Supplemental Figure 4.**
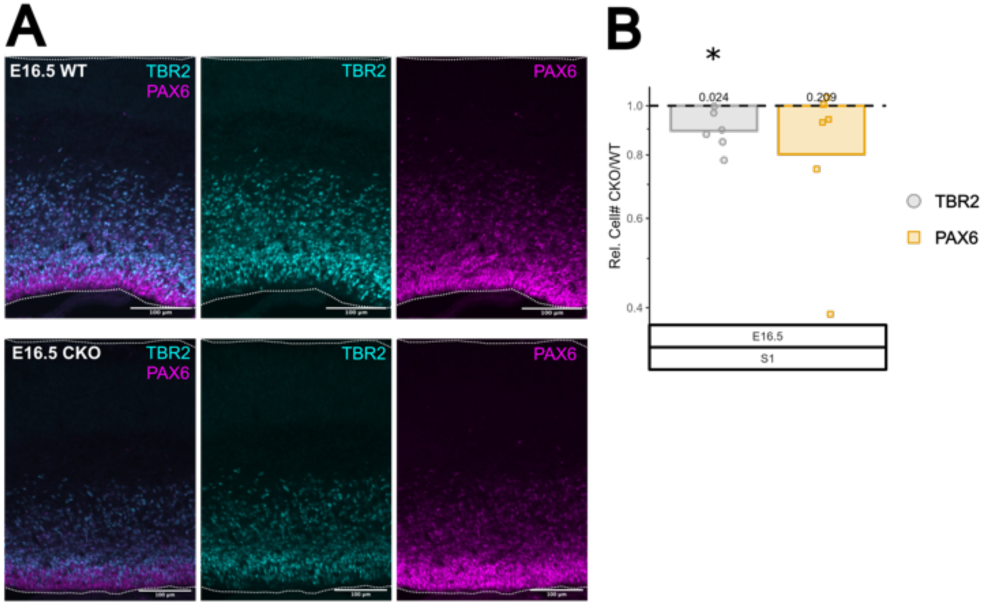
Intermediate progenitor cells are reduced at E16.5 in *vGluT2*cKO cortex. **A.** Immunostaining of littermate WT (left) and *vGluT2* cKO (right) sections with TBR2 (intermediate progenitor cells) and PAX6 (radial glia) at E16.5. **B.** Analysis of relative cell number for progenitor markers (TBR2 and PAX6) at E16.5, showing a reduction in TBR2^+^ IPCs and no change in PAX6^+^ radial glia. Data are shown as the relative difference of *vGluT2* cKO to WT littermates (dashed line). Points represent individual pairs and bars represent pooled mean. *P < 0.05, **P < 0.01, ***P < 0.001.

